# Pan-Prediction of MHC-II Restricted Epitopes Across Species via an Alphafold-based Quantification Scheme

**DOI:** 10.1101/2024.10.11.617946

**Authors:** Suqiu Wang, Lingming Kong, Dongmei Hu, Liangzhen Zheng, Caiyi Fei, Liubao Du, Ziche Tang, Sheng Wang, Shi Xu, Hanchun Yang, Nianzhi Zhang

## Abstract

Predicting MHC-II restricted epitopes across species used to be challenging, but Alphafold (AF) may provide a structure-based pan-prediction solution. In this study, we established the new tool AF-pred with a clear standard for quantitative prediction results. Compared to the sequence-based tools heavily trained with human ligandome, AF-pred does not show advantage in predicting the binding patterns of human HLA-II but has far better performance in predicting binding patterns of other animals’ MHC-II. Using recently resolved bat MHC-II structures, we analyzed AF-pred’s prediction capability, logic and limitation. In addition, we also explored the impact of AF algorithm iterations on the prediction of MHC-II restricted epitopes. The results demonstrated that AF-pred is capable of cross-species prediction of MHC-II restricted epitopes and is conducive to the development of novel veterinary vaccines.

## Introduction

Control of animal infectious diseases, especially zoonoses, is very important for both livestock industry and public health^1,2^. Developing safe and effective veterinary vaccines is the most effective approach for disease control^3,4^. However, one of the key challenges for vaccine design is determining effective T cell epitopes in pathogen proteomes^5–7^. T cell epitopes are peptide ligands bound to Major-Histocompatibility Complex (MHC). They are further divided into endogenous epitope peptides presented by MHC-I, and exogenous epitope peptides presented by MHC-II^8^. The evolution of animals’ immune system led to extremely high diversity of MHC: the MHC protein sequences show significant variation across species^9–11^; and even within human genome, MHC shows amazing polymorphism: as of the recent IMGT/HLA database, there are over 29,000 known human MHC alleles identified, including both MHC-I and MHC-II (https://www.ebi.ac.uk/ipd/imgt/hla/index.html). Since incorporating T cell epitopes into vaccines is crucial for increasing vaccines’ antigenicity, it would be ideal to have a revolutionary tool for predicting MHC-restricted T cell epitopes across various species to overcome the MHC diversity. The prediction of MHC-restricted epitopes relies on the understanding of the structure of peptide-MHC complexes (pMHC) and the analysis of MHC ligandome^12–14^. The extracellular regions of MHC-I and MHC-II have comparable peptide-binding grooves composed of a β-sheet topped by two antiparallel α-helices. Some inherent positions in the peptide-binding grooves are known as pockets, which can accommodate the side chains of amino acid residues at specific positions in peptides and play an important role in anchoring peptides. Statistical analysis of MHC ligandome reveals that specific positions in peptide ligands have obvious preferences on certain amino acid residues, and such rules are determined by the properties of the corresponding pockets^15–17^. Due to differences in structure and peptide binding characteristics, the prediction of MHC-II restricted epitopes is more challenging than that of MHC-I epitopes^18^.

Regarding to the structure, MHC-I consists of a heavy chain (containing α1-α3 domains) and a light chain (β2m). β2m does not have polymorphism except in fishes, and the peptide-binding groove of MHC-I are determined by the α1 and α2 regions of the heavy chain^19^. However, MHC-II is a heterodimer composed of an α chain (containing α1 and α2 domains) and a β chain (containing β1 and β2 domains), and the peptide-binding region is formed by the α1 and β1 domains. The polymorphic α and β chains bring overwhelming combinations and diversity to the peptide-binding grooves^20^.

Regarding to the peptide binding characteristics, most MHC-I have peptide-binding grooves closed at both ends that participate in fixing the peptide ligand. However, the peptide-binding groove of MHC-II molecules is open, allowing the peptide ligand to extend out from both ends of the groove^21^. This leads to significant differences in MHC-I and MHC-II ligandomes: MHC-I peptide ligands usually have 8-12 amino acids (aa), typically 9aa; while MHC-II peptide ligands are much longer, usually 12-25aa. Actually, the core binding region in the MHC-II peptide-binding groove is also 9aa^15^. Furthermore, MHC-II binding peptides exist in two possible orientations: forward and reverse^22^. Therefore, accurate identification of the 9aa-binding core region of each peptide ligand is crucial for in-depth analysis of MHC-II peptide binding motif—but this is not an easy task.

Moreover, among the 6 pockets (A-F) of the MHC-I peptide-binding groove, pockets B and F are known to play the major role, making the peptide binding motif of MHC-Irelatively clear. It is usually manifested as the preference of pockets B and F for the anchoring residues at the second position (P2) and the C-terminal (PΩ). However, in the peptide-binding groove of MHC-II, 4 pockets bind to the residues at positions P1, P4, P6, and P9 in the 9aa-binding core region, respectively^15^. This obviously contributes to the complexity of MHC-II peptide binding motif. So far, various tools for predicting MHC-restricted epitopes roughly fall into 2 categories^12,13,23–25^: the tools based on deep learning of MHC ligandome sequences only (not including any structure biology information), and the tools based on the structure prediction of pMHC complexes. Sequence-based tools have far better performance in predicting MHC-I restricted epitopes than MHC-II restricted epitopes^26^, and such difference is actually determined by deep learning model’s requirement of massive training data: if the data feed is below a certain threshold, the trained model will show poor prediction capability^27^.

In public database like IEDB.org, known MHC-II ligands are much fewer than MHC-I ligands. This is because eluting MHC-II ligands is usally more challenging in wet lab: MHC-I is universally present on the surface of nucleated cells, whereas MHC-II is mainly located on the surface of antigen-presenting cells^28^, which limited the sources of obtaining MHC-II ligandome.

In recent years, the accumulation of mass spectrometry data of MHC-II ligandome and the development of sequence-based tools have improved the prediction capability of MHC-II restricted epitopes^26,29,30^. However, these mass spectrometry data and prediction tools are mainly compatible for human MHC-II (HLA-II) molecules. Although certain tools like NetMHCpan can perform pan-prediction based on the similarity of aa compositions in the MHC binding groove, they have poor performance in predicting non-human MHC-II restricted epitopes^27^. MixMHC2pred has already involved MHC-II ligandomes of mice, cattle, and chickens for model training, but they are still insufficient for feeding the deep-learning model that requires massive sequence data^31^. Since animal MHC-II ligandome data are unlikely to very rapidly increase in the near future, prediction models based on deep-learning models might all reach an upper limit that is not high. In such a situation, the tools based on the structure prediction of pMHC complexes would be a good alternative, because they don’t require massive training data to feed the model.

Fortunately, AlphaFold2 (AF2) with greatly improved protein structure prediction accuracy and credibility are now providing the foundation for developing new tools based on the structure prediction of pMHC complexes^32^. Some studies that applied AF2 to HLA structure analysis achieved good prediction accuracy as well as better interpretability for the prediction results^33–35^. Our research on bat MHC-I molecules has proven that the pMHC structure predicted by AF2 multimer (AF2M) closely mimics the real crystal structure, indicating that AF2M can also be used for the prediction of non-human MHC epitopes^36,37^. Recently, the new AF3 algorithm has further improved capability of predicting protein complex structures^38^, and this gave us the chance to develop cross-species MHC-II restricted epitope prediction methods based on AF (AF2M/AF3) structure prediction and wet-lab data verification.

In this study, based on the credibility of AF-predicted protein structures, we developed the structure-based tool AF-pred for predicting MHC-II restricted epitopes in a quantitative approach. We have validated AF-pred with peptide binding data and resolved crystal structures of MHC-II molecules from humans, pigs, cattle, and bats. Overall, for MHC-II of human and other species, the epitopes predicted by AF-pred show high level of consistency with wet-lab data. Moreover, the iteration and improvement of the AlphaFold series structure prediction tools also significantly contribute to accurate prediction of MHC-II restricted epitopes. AF3’s prediction can better reflect the factors restricting peptide binding, with higher accuracy and better interpretability of the prediction results; while AF2M can provide diverse prediction results that are helpful for discovering various peptides binding modes of MHC-II.

Compared with the resolved crystal structures, AF-pred’s predictions can almost cover all possible interactions between peptide ligands and MHC-II peptide-binding groove. However, its preference on the most stable binding mode with the conventional peptide conformation may lead to neglection of unconventional binding modes. Compared with sequence-based tools that really look like black-boxes, AF-pred has natural advantages in prediction result interpretability and evaluation of peptide binding forces. Our research demonstrated successful application of AF in predicting pMHC-II structures, and AF-pred is a feasible solution for cross-species prediction of MHC-II restricted epitopes. This will greatly promote immunology research in various animal species and facilitate the design of veterinary vaccines.

## Results

### Development of the AF-pred Prediction Tool for MHC-II restricted epitopes and Validation based on HLA-II ligandome

Accurate identification of the 9aa-binding core region of the MHC-II peptide ligand has always been very challenging in the development of sequence-based prediction tools that direct convert peptide sequences into vectors via coding^12,13^. However, from the perspective of structure biology, accurate identification of the 9aa-binding core region is a question that can be solved in a generalized method. The closed binding groove of MHC-I leads to highly diverse conformations in the middle part of the peptide ligand, whereas the open binding groove and pocket distribution of MHC-II enable the peptide ligand to fully extend, presenting a conserved conformation akin to the polyproline type II helix (Fig. 1a). In the 9aa-binding core region of the MHC-II peptide ligand, the relative spatial positions of the Cα atoms of each residue within the binding groove are extremely conserved, with deviation amplitude typically <2.5 Å. This conserved peptide binding characteristic provides a distinct criterion for evaluating prediction results, and it is very helpful for developing an epitope prediction tool based on the structure prediction of pMHC-II complex. AF2M has notably enhanced credibility of complex structure prediction validated by pMHC complex structures^37^. Moreover, the latest AF3 has further augmented the capability of predicting complex structures^38^.

**Figure 1.**
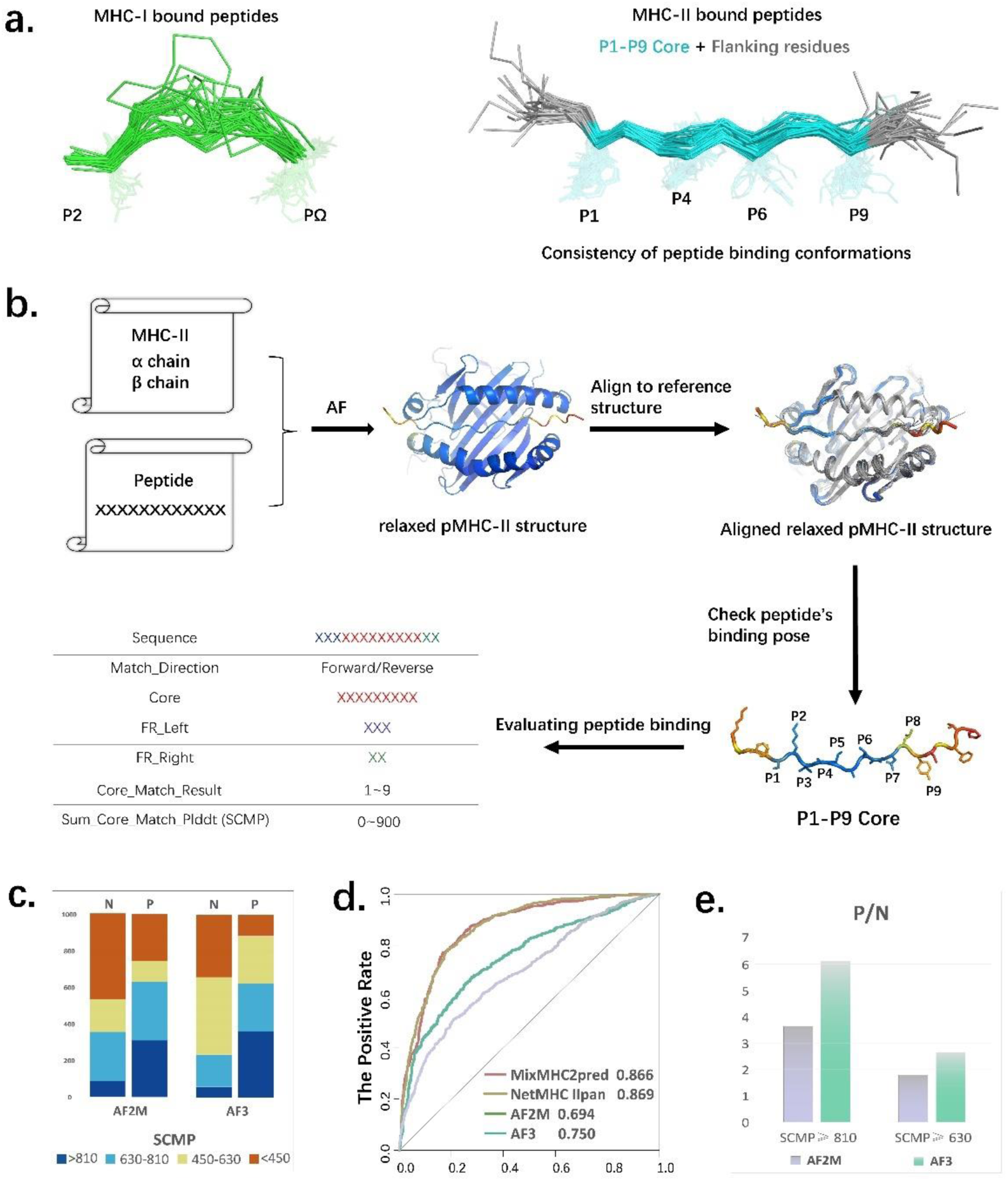
AF-pred: a prediction scheme for MHC-II binding peptides based on the prediction of the AlphaFold, along with its validation. **(a)**. Comparison of the binding conformations between MHC-I and peptides versus MHC-II and peptides. **(b)**. Flowchart of the prediction and evaluation scheme for MHC-II binding peptides. Validation of AF-pred using AF2M and AF3 on public datasets, illustrated by SCMP distribution **(c)**, ROC_AUC scores **(d)** and P/N ratios **(e)**.

On this basis, we developed a novel MHC-II eitope prediction tool AF-pred, as illustrated in Fig. 1b: The optimal predicted model of the target pMHC-II complex was selected, and then it was automatically compared with the resolved pMHC-II complex structure in reference. With 2.5Å as the threshold of the spatial distances of the Cα atoms from the counterparts in reference peptide ligands, the corresponding P1-P9 residues in the predicted peptide (Core_Match_Result) were determined, along with the binding orientation and flanking regions (FRs) of the peptide. Finally, the sum of pLDDT values of the spatially matched Cα atoms of the predicted peptide (Sum_Core_Match_Result, SCMP) is calculated for reflecting the probability of peptide binding. The SCMP scores are divided into 4 intervals: > 810 (high confidence), 630-810 (medium confidence), 450-629 (low confidence), and <450 (very low confidence). High SCMP scores of the predicted peptide generally indicate high probability of binding to the target MHC-II molecule. In summary, the prediction capability of AF-pred derive from two aspects: the credibility of AF prediction and the conservation of pMHC-II conformation.

To validate the prediction capability of AF-pred, 1000 positive (binding) peptides and 1000 negative (non-binding) peptides from publicly available HLA-II datasets were selected, each from 10 different allele combinations (Supplementary Table 2). Subsequently, they were scored using AF-pred based on AF2M and AF3 (AF2M-pred and AF3-pred), and the SCMP score distribution was illustrated in Fig. 1c. Regarding to SCMP score distribution, the positive peptide set have generally higher SCMP scores than the negative peptide set; moreover, such difference is more significant in the prediction results of AF3-pred than those of AF2M-pred, indicating AF3-pred may have superiority in distinguishing binding and non-binding peptides. After ROS scoring of this data, the AUC area of AF3-pred is 0.75, which represents a significant improvement over the 0.694 of AF2M-pred, reaching the application standard. Regarding to the AUC, AF-pred still has a gap from the two sequence-based tools, NetMHCIIpan^26^ and MixMHC2pred^31^. This is probably because they have been heavily trained with massive HLA-II ligandome data. However, the major advantage of AF-pred is its prediction capability not relying on massive ligandome data, which is especially useful in the fields with very limited ligandome data.

Predictive tools typically have scoring methods to determine the confidence of results. Consequently, the accuracy of high-score results reflects the tools’ credibility. Among the high confidence peptides with SCMP scores > 810, the P/N value of AF3 was about 6.1, while that of AF2M was about 3.6. Among the medium confidence peptides with SCMP scores between 630 and 810, the P/N values of AF3 and AF2M were about 2.8 and 1.8, respectively (Fig. 1d). The test results above validated the AF-pred’s capability of screening peptide ligands that bind to MHC-II, as well as the strong positive correlation between the SCMP score and the probability of actual binding.

### Validation of AF-pred’s cross-species prediction capability for MHC-II restricted epitopes

For predicting human HLA-II peptide ligands, AF-pred is less competitive than NetMHCIIpan and MixMHC2pred heavily trained with corresponding ligandome data. However, for predicting peptide ligands of animal MHC-II whose ligandome data is very limited, AF-pred based on structure biology elements may transcend species difference and the diversity of MHC-II heterodimer combinations. MixMHC2pred includes some bovine and chicken MHC-II ligandome data in its training set, so it could provide certain cross-species predictions. Therefore, we used MixMHC2pred as a reference tool for head-to-head comparison with AF-pred^31^. We conducted the prediction on porcine MHC-II (SLA-DRA*01:01:01_DRB*01:01:01) with the African swine fever virus (ASFV) P72 protein (Fig. 2a) and bovine MHC-II (BoLA-DRA*01:01_DRB3*16:01) with the foot-and-mouth disease virus (FMDV) polyprotein (Fig. 2b). The prediction results from AF2M-pred, AF3-pred, and MixMHC2pred exhibited significant differences. Among the peptides with high ranking by these 3 prediction tools, the predictions of their 9aa-binding core regions are not completely consistent. Among 18 peptides, only 9 peptides have completely identical 9aa-binding core regions predicted by 3 tools. Even for the same peptide, AF2M-pred and AF3-pred may give different predictions, such as peptides P185 and P417 in the ASFV P72 protein, and peptides P553, P657, P793, and P2017 in the FMDV. Notably, AF3-pred predicted that peptide P417 may bind to SLA-DRA*01:01:01_DRB*01:01:01 in a reverse orientation. Moreover, even if the 3 tools have consistent prediction for the 9aa-binding core region of a peptide, they may give quite different scores. Peptides P233, P289, and P481 in the P72 protein, and peptides P57 and P1321 in the FMDV are examples. The differences in prediction results between AF-pred and MixMHC2pred reflect the differences between sequence-based and structure-based prediction schemes, while the differences in prediction results between AF2M-pred and AF3-pred reflect the impact of AF algorithm iterations. *In-vitro* binding assays were conducted to validate these peptides: 4 top-ranked peptides predicted by MixMHC2pred were false-positive (non-binding), whereas only one high-score peptide each from AF2M-pred and AF3-pred was false-positive, suggesting the superior prediction capability of AF-pred for cross-species MHC-II restricted epitopes.

**Figure 2.**
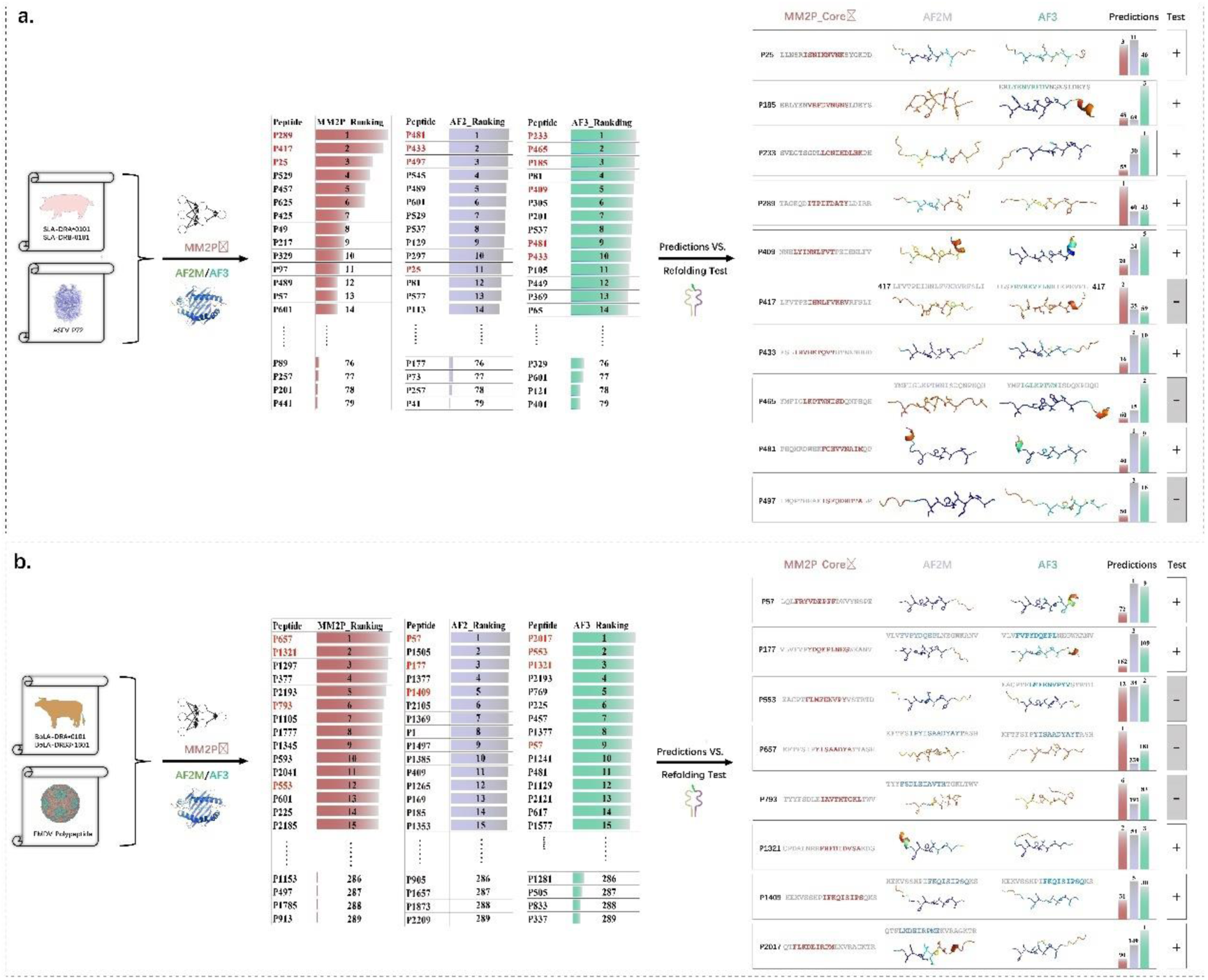
Validation of AF-pred’s cross-species capability in predicting MHC-II binding peptides. Comparative tests using porcine MHC-II with African swine fever virus P72 protein **(a)** and bovine MHC-II with foot-and-mouth disease virus protein **(b)**. Results indicate significant differences among the predictions from AF2M-pred, AF3-pred, and MixMHC2pred, suggesting distinct performances across these prediction methods when applied to different species and viral proteins. The peptides were ranked by their scores predicted by the three tools, respectively. The bars represent the scores of the predicted peptides. The peptides with high score were selected and tested by *in vitro* refolding.

To further validate the cross-species prediction capability of AF-pred for MHC-II restricted epitopes, we used bat MHC-II (EF-DRA and EF-DRB1) and an overlapping peptide library covering the RBD region of the COVID19-S protein for the test. AF2M-pred and AF3-pred each predicted 7 peptides with SCMP scores greater than 630 with more than medium confidence. Among these 4 peptides (P127, P169, P225, and P295) were consistent in their prediction results (Fig. 3a). *In-vitro* binding experiments showed that all predicted peptides with more than medium confidence bind to Bat-MHC-II_EF (the blue bars represent the peak of the refolded complexes purified by gel filtration). Moreover, for the 4 overlapping peptides consistent in AF2M-pred and AF3-pred prediction results, AF2M-pred predicted 3 high credibility peptides (P127, P225, and P295), while AF3-pred predicted only 1 high credibility peptide (P169). The differences in SCMP scores of these 4 peptides mainly originated from the anchor residues at positions P4 and P9 (Fig. 3b). The predictions of the peptide-binding groove of Bat-MHC-II_EF by AF2M-pred and AF3-pred were very consistent, both showing that the P4 pocket is negatively charged, while the P9 pocket is positively charged. In terms of the rationality and interpretability of the anchor residues and pocket, the results of AF3-pred are more reasonable, especially for peptide P169. The 9aa-binding core regions of P169 predicted by AF2M-pred and AF3-pred are inconsistent. Obviously, the P9-D predicted by AF3-pred is more suitable for the positively charged P9 pocket. For the other two peptides, P225 (ISTE<u>IYQAGSTPC</u>NGVEG) and P295 (LSFE<u>LLHAPATVC</u>GPKKS), the predicted P9 anchor residues are both uncharged cysteine, which does not match the positively charged P9 pocket. AF3-pred gave them poor credibility, but AF2M-pred believed they bind well.

**Figure 3.**
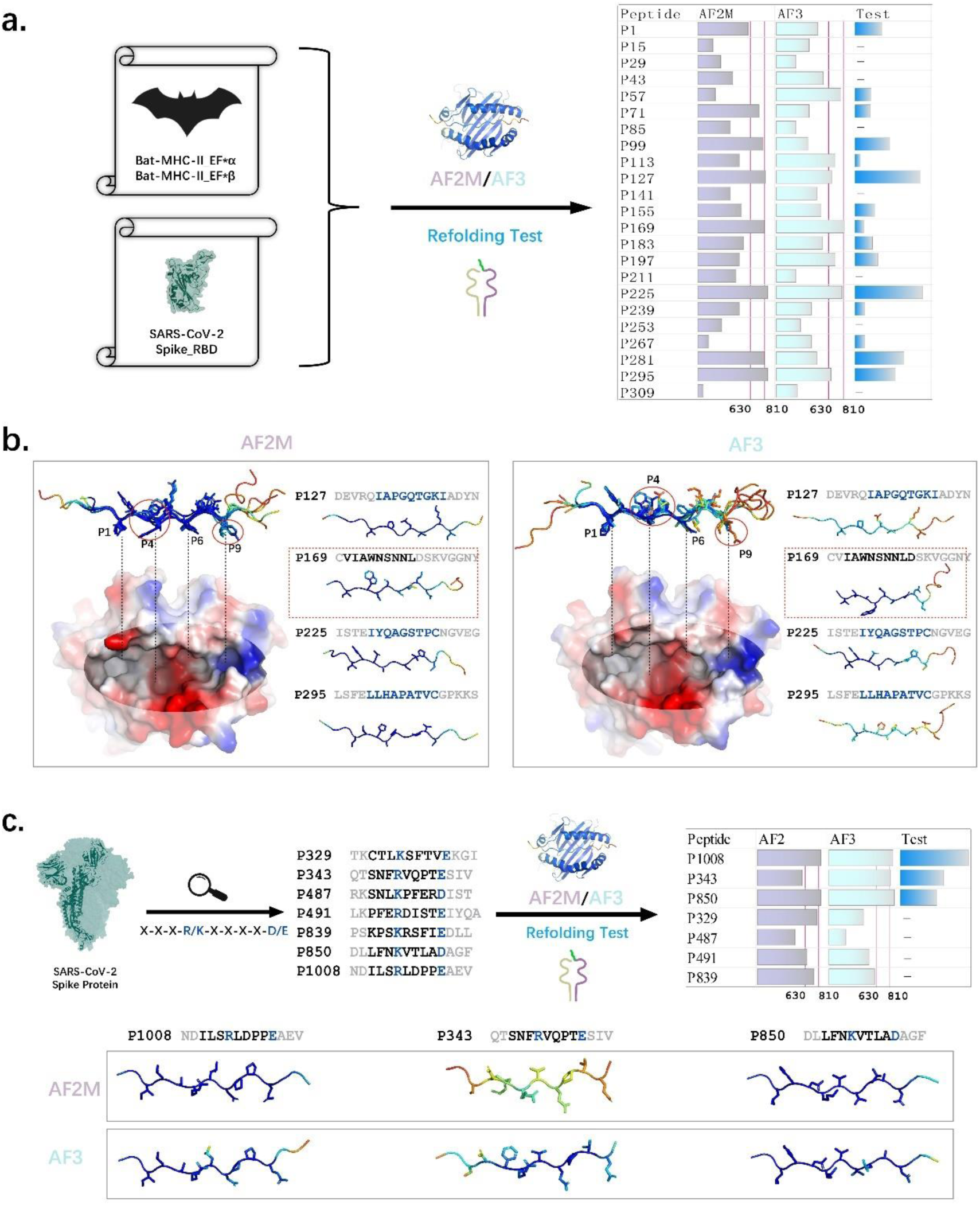
Validation of AF-pred’s ability to predict MHC-II binding peptides in bats. **(a)**. Testing with Bat-MHC-II_EF and an overlapping peptide library covering the RBD region of the COVID-19-S protein. The performance of AF-pred was validated by the *in vitro* refolding. **(b)**. Comparative analysis of the 4 overlapping peptides consistent in AF2M-pred and AF3-pred prediction results. A detailed examination of the predicted conformations reveals notable differences primarily centered around the anchor residues at positions P4 and P9. **(c)**. Further validation the suitability of AF-pred demonstrate that under conditions where anchor residues align optimally with MHC pockets, AF3-pred exhibits superior accuracy.

We speculated that AF3-pred pays more attention to the constraints for the complex formation, making results more interpretable. In contrast, AF2M-pred is not sensitive to the restriction factors for complex formation, leading to more chaotic prediction results that may be difficult to interpret. Bat-MHC-II_EF can bind to 15 of the 23 tested peptides, indicating that its peptide binding motif is not very restrictive, so it may be more favorable to AF2M-pred. For more clearly defined interaction patterns, the prediction performance of AF3 is better. To verify our speculation, total 7 peptides in the S protein that match the charge properties of the P4 and P9 pockets of Bat-MHC-II_EF were selected to run the test (Fig. 3c). The prediction results of AF3-pred were completely consistent with the wet lab result that 3 of the 7 peptides (P343, P850, and P1008) bind to Bat-MHC-II_EF to form complexes. However, AF2M-pred’s SCMP score for P343 failed to reach the credibility threshold of 630. Moreover, AF2M-pred gave higher SCMP scores to the other 4 negative peptides than AF3-pred, and 3 of them are above the credibility threshold of 630. The better performance of AF3 in this test indicated that AF3 may be more suitable for prediction tasks with clear restrictive interaction factors.

### Comparison with the novel pBat-MHC-II crystal structure reveals the logic and capability boundaries of AF-pred

The comparison of predicted pMHC-II structure with resolved crystal structures provide fundamental assessment of AF-pred’s capabilities. We have elucidated the crystal structure of the complex between the P1008 peptide from the COVID19-S protein and Bat-MHC-II_EF (designated as pBat_EF-NDIL), with detail crystallographic data presented in Table 1. The unit cell of pBat_EF-NDIL crystal structure contains two molecules (M1 and M2). Unexpectedly, the bound peptide conformations in M1 and M2 are not entirely consistent, yet clear electron density maps indicate that both conformations are stable (Fig. 4a). The differences mainly occur at the P8-P9 positions, revealing that even a single peptide could have more than one conformations compatible for binding MHC-II. The peptide conformations predicted by AF2M-pred and AF3-pred are consistent with that in M2, which represents the canonical MHC-II peptide binding conformation (Fig. 4b). For the conserved interactions between the peptide main chain and the MHC-II peptide-binding groove, the prediction of AF-pred is highly consistent with the real structure of pBat_EF-NDIL (red marked residues), except the hydrogen bond between the main chain of the P8 residue and β61W. This bond predicted by both AF2M-pred and AF3-pred is common in other resolved pMHC-II structures, but actually does not exist in M1 and M2. The interactions of the peptide side chain in M1 and M2 are not completely the same. For the hydrogen bonds that exist in both M1 and M2, AF2M-pred did not predict the hydrogen bond between P4-R and β26Y, while AF3-pred did not predict the hydrogen bond between P4-R and β74S. However, for both AF2M-pred and AF3-pred, the total predicted hydrogen bonds and salt bridges between the peptide and EF are more than those in the M1 and M2 conformations. The above implies that AF-pred prefer to give peptide conformation with more possible interactions within a reasonable range, but it cannot predict non-classical MHC-II peptide binding conformations.

**Figure 4.**
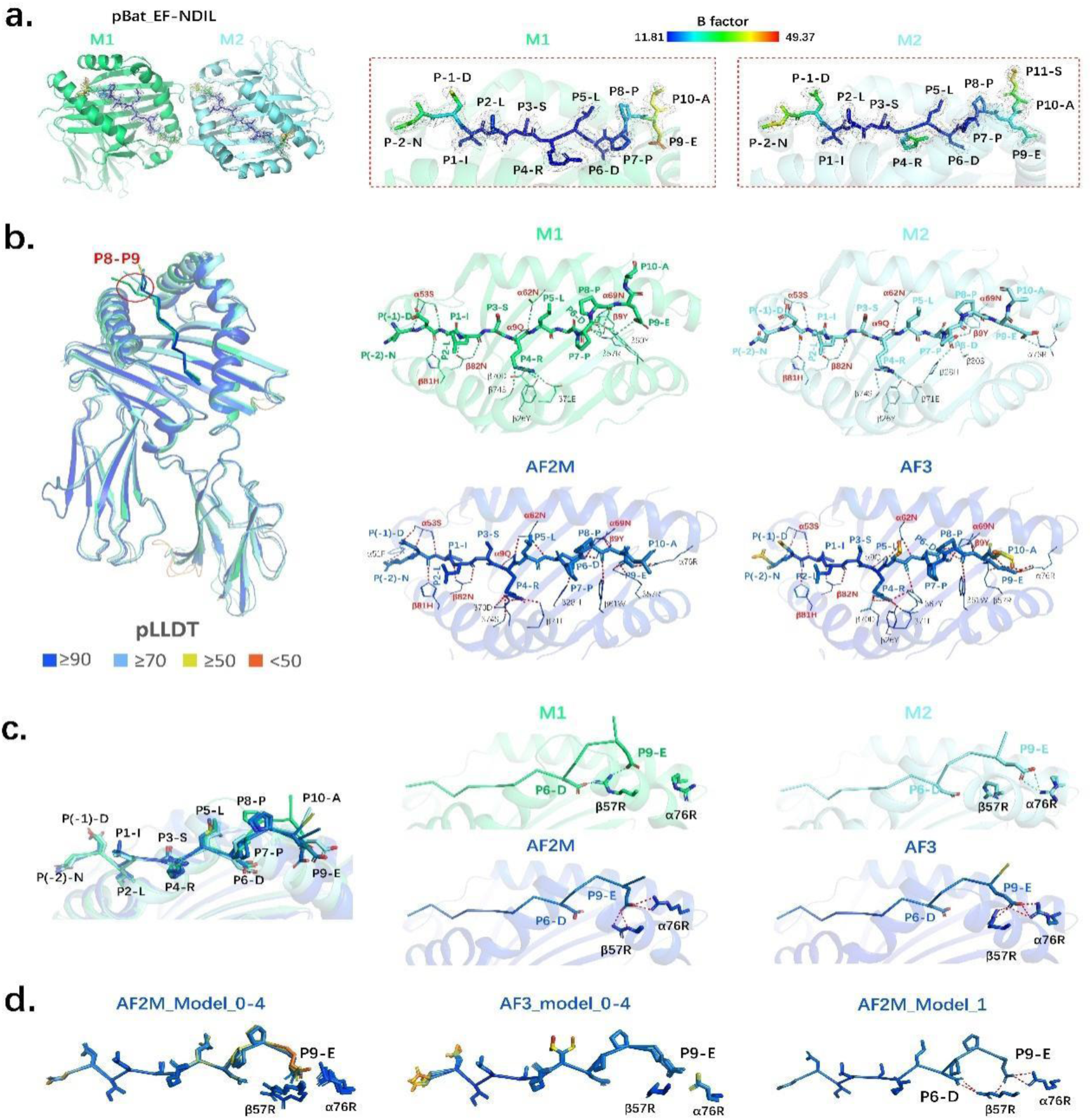
Comparison of MHC-II peptide binding conformations predicted by AF models versus crystallographic structures. **(a)**. The crystallographic structure of the pBat_EF-NDIL complex, featuring the P1008 peptide from the COVID19-S protein bound to Bat-MHC-II_EF, was resolved, revealing two distinct conformations: M1 and M2. **(b)**. Predicted structures from AF2M and AF3 closely resemble the peptide conformation in M2. The prediction of AF-pred is highly consistent with the real structure of pBat_EF-NDIL for the interactions between the peptide and the conserved MHC-II residues (red marked). **(c)**. Conformational variations in peptides between the pBat_EF-NDIL crystal structure and predictions are primarily attributed to the flexible α76R and β57R. **(d)**. Comparation of predicting structures generated by AF2M versus AF3 reveals that AF2M exhibits greater diversity in its predictions.

**Table 1.**
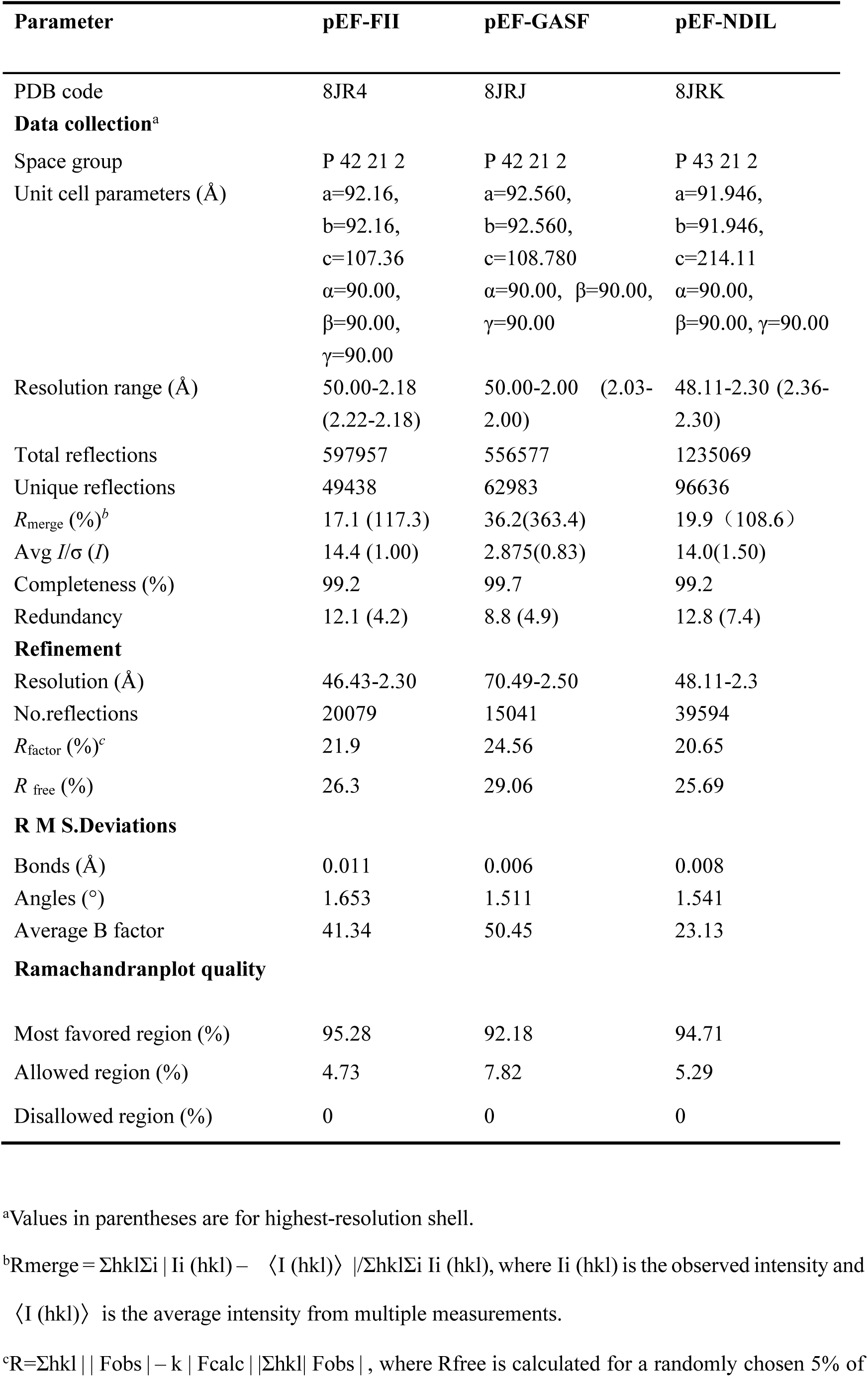
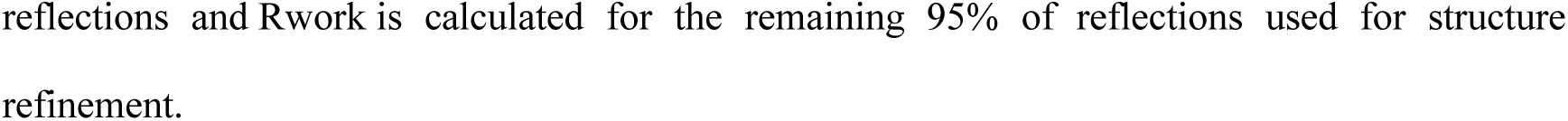
X-ray diffraction data processing and refinement statistics.

The disparity in peptide conformation predominantly originate from the highly flexible and positively charged arginine residues, α76R and β57R, within the EF molecule (Fig. 4c). In conformation M1, β57R engages in interactions with both P6-D and P9-E, whereas α76R does not establish a salt bridge with P9-E, leading to an arched backbone of the P6-P9 peptide segment in M1. Conversely, in M2, β57R is not involved in binding, with only α76R forming a salt bridge with P9-E. The predicted structures suggest that both α76R and β57R concurrently bind to P9-E, resulting in an increased number of salt bridges. The AF methodology is capable of producing multiple predictive structures for an individual molecule, typically presenting 5 optimal configurations (Models 0 to 4). Only the Model_0 is used for the above analysis. This led to an inquiry into AF-pred’s capability to forecast variable binding conformations (Fig. 4d). Upon comparison of the quintet of predicted structures, it became evident that AF2M-pred could anticipate alterations involving α76R and β57R, whereas in the multiple outputs from AF3-pred, the binding modes for these residues remained invariant. Specifically, AF2M_model_1 could foresee β57R’s concomitant engagement with P6-D and P9-E, with α76R also implicated in the binding at P9-E. These findings indicate that AF2M-pred may provide more diversified predicted structures than AF3-pred, and this feature is useful for delineating diverse peptide binding conformations.

### Successful modification of peptide ligands based on AF-pred prediction is the proof of the accuracy

Both AF2M-pred and AF3-pred predicted that the peptide (PGDSIIRSMPEQTSEK) derived from the crystallographic complex of avian MHC-II (BL2*019:01) could bind Bat-MHC-II_EF, and the 9aa-binding core region is IRSMPEQTS. Notably, in contrast to AF3-pred, AF2M-pred gave an alternative result of 9aa-binding core region (DIIRSMPEQ) in Model_1, but the P1-D residue could not be accommodated within the P1 pocket (Fig. 5a). Upon mutating P1-D to P1-F, AF2M-pred provided the prediction that both IRSMPEQTS and <u>F</u>IIRSMPEQ could act as the 9aa-binding core region, but the SCMP score of <u>F</u>IIRSMPEQ is lower than 630. In the contrast, AF3-pred regarded <u>F</u>IIRSMPEQ as the exclusive candidate of 9aa-binding core region. For the peptide sequence FIIRSMPEQ, both AF2M and AF3 anticipated diminished credibility at the C-terminus. Given that the P9 pocket of Bat-MHC-II_EF is biased towards binding negatively charged residues, we introduced the second mutation by inverting the positions of P8-E and P9-Q to FIIRSMP<u>QE</u>, making P9-E more compatible with the positively charged P9 pocket. Upon the second mutation, AF2M-pred indicated that both FIIRSMP<u>QE</u> and IRSMP<u>QE</u>TS could serve as the 9aa-binding core region, and FIIRSMP<u>QE has higher</u> SCMP score. However, AF3-pred predicted FIIRSMP<u>QE</u> as the only viable 9aa-binding core region with enhanced SCMP score. *In-vitro* binding assays for these peptides confirmed the binding of both the native and modified peptides to Bat-MHC-II_EF. Thermodynamic stability measurements of the complexes revealed a significant increase in binding affinity following the second mutation, implying a possible alteration in the binding pattern of the mutated peptide relative to the native peptide.

**Figure 5.**
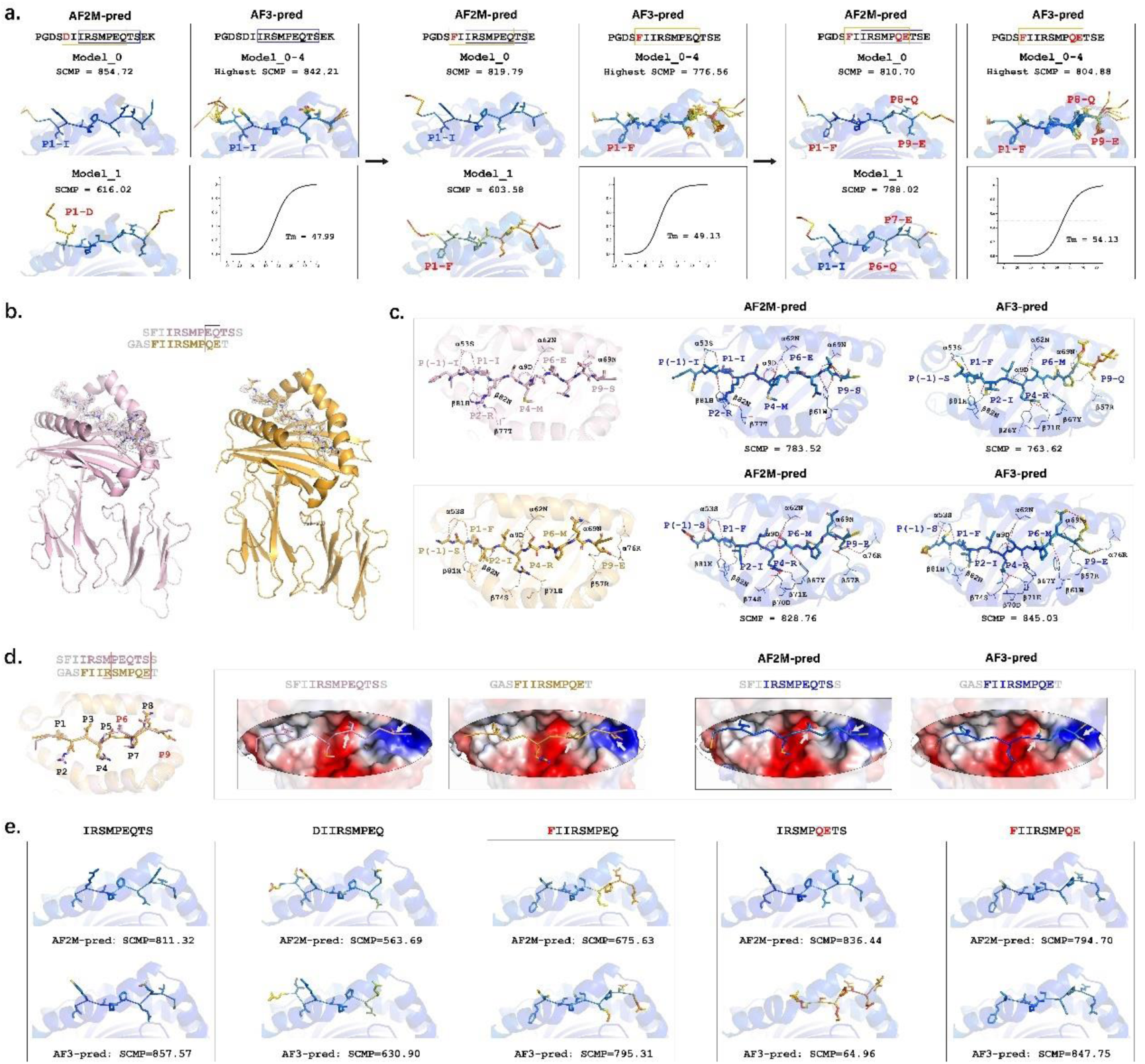
Modification of peptide ligands based on AF-pred prediction. **(a)** Predictive analysis of peptide PGDSIIRSMPEQTSEK and its mutant sequences with Bat-MHC-II_EF by AF-pred. AF2M-pred is capable of find more possible 9aa binding core regions. **(b)** The electron density maps of the 9aa binding core regions (colored) and the part flanking regions (gray) of the peptides. **(c)** Comparative analysis of crystal structures of pBat_EF-PGDS and pBat_EF-GASF and their AF-pred predications. **(d)** Highlighting differences in peptide conformations at position P6. **(e)**. Predictions of possible 9aa binding core regions in the peptide PGDSIIRSMPEQTSEK and its mutants that bind to Bat-MHC-II_EF.

While retaining the 9aa-binding core region, after multiple modifications of the peptide FR guided by AF-pred, we ultimately resolved the complex crystal structures of two mutant peptides (PGDS<u>F</u>IIRSMPEQTS and G<u>A</u>S<u>F</u>IIRSMP<u>QE</u>TS) in conjunction with Bat-MHC-II_EF (referred as pBat_EF-PGDS and pBat_EF-GASF, respectively), with detailed crystallographic data delineated in Table 1. Because the ends of these two peptides extend outside the binding groove, some of those residues lack electron density maps. However, the electron density maps of their respective 9aa-binding core regions are complete and clear. (Fig. 5b). In contrast to pBat_EF-NDIL, the crystal structure of pBat_EF-GASF illustrates that P9-E is capable of engaging in simultaneous salt bridge formations with both α76R and β57R within the EF domain, and this result is consistent with the prediction of AF-pred (Fig. 5c). This substantiates the proficiency of AF-pred in exploring potential structural interactions and peptide ligand optimization based on more stable conformation.

Furthermore, it is noteworthy that the differences in the peptide ligand conformations between pBat_EF-PGDS and pBat_EF-GASF are most evident at the P6 position (Fig. 5d). In pBat_EF-PGDS, P6-E is embedded into the designated pocket, with its side chain towards the charged polar region within the pocket; whereas the short and uncharged P9-S side chain, naturally extends downward. In pBat_EF-GASF, the hydrophobic side chain of P6-M matches the hydrophobic region of the pocket, and P9-E with a negative charge extends laterally to form a salt bridge with the corresponding arginine residue. This reveals the flexibility of the Bat-MHC-II_EF binding pocket to interact with anchor residues of diverse characteristics and form rational binding conformations. AF-pred’s sophisticated exploration on these intricate binding conformations highlights its prediction capability.

The crystallography evidence indicates that the 9aa-binding core regions of all the native and mutant peptides are consistent with the predictions of AF2M-pred. Notably, AF2M-pred can discover two different possible 9aa-binding core regions in a peptide and rank them correctly (Fig. 5a and c). In contrast to AF2M-pred, the top-ranked structures predicted by AF3-pred did not provide such desirable diversity, even though multiple runs were performed. This suggests that the prediction mechanism of AF2M-pred may have more stochastic elements than that of AF3-pred. From this perspective, AF2M-pred might have certain advantage in exploring a broader spectrum of peptide binding conformations.

For AF3-pred, it is meaningful to explore the logic behind its predictions of the native and first-round mutant peptide (PG<u>F</u>SIIRSMPEQTSEK). For the native peptide, AF3-pred gave a high-confidence SCMP score to the 9aa-binding core region IRSMPEQTS (Fig. 5a). Conversely, for the mutant peptide PG<u>F</u>SIIRSMPEQTSEK, the predicted 9aa-binding core region shifted from IRSMPEQTS to FIIRSMPEQ, with enhanced restrictive binding of P4-R, notwithstanding a notable reduction in the credibility assigned to P6 and P9 (Fig. 5a and c). These observations reinforce our hypothesis that AF3-pred could more effectively identify restrictive factors for peptid-MHC-II binding.

Even the flanking regions of the peptide can influence AF3-pred’s output of the 9aa-binding core region. For example, the SCMP score of the 9aa-binding core region <u>F</u>IIRSMP<u>QE</u> in peptide PGDS<u>F</u>IRSMP<u>QE</u>TSE is 804.88 (Fig. 5a). However, the reduction and fine-tuning of the N-terminal and C-terminal flanking regions led to enhanced SCMP score of <u>F</u>IIRSMP<u>QE</u> in G<u>A</u>S<u>F</u>IIRSMP<u>QE</u>TS (Fig. 5c). Predicting the binding of the isolated 9aa-binding core region to MHC-II may provide a “cleaner” output. When the flanking regions were excluded, the AF3-pred SCMP scores would be ranked as IRSMPEQTS > DIIRSMPEQ, IRSMPEQTS > FIIRSMPEQ, and FIIRSMPQE > IRSMPQETS (Fig. 5e). The results above are consistent with the crystal structure. Notably, AF3-pred gave a higher SCMP score to IRSMPEQTS than to FIIRSMPEQ, which contrasts with the outcomes from the first-round mutated peptides (Fig. 5a). This indicated that the labile nature of the peptide FRs increase the difficulty of prediction. More focused prediction of the 9aa-binding core region regardless of FRs is reasonable simplification that not only reduce the computation complexity but also lead to more accurate prediction results.

### AF-pred is not capable of predicting non-canonical binding mode of peptide and MHC-II

Although the conformation of peptide ligands bound to MHC-II is typically conserved, it is well-known that certain MHC-II peptide ligands exhibit non-canonical binding modes. In our study, AF-pred showed insufficient capability of exploring such non-canonical binding modes. We intentionally selected certain elucidated pMHC-II structure with non-canonical binding modes as the teststone (Fig. 6a). Among the predicted structures of these pMHC-II complexes provided by AF-pred, with the exception of the PDB entry 3LQZ, all the other predicted structures were not consistent with the crystal structures (Fig. 6b). The peptide binding conformations were determined either low confidence (PDB entries 6T3Y and IUVQ) or mistakenly complying with the canonical binding modes. For example, the avian MHC-II molecule BL2*02 (PDB entry 6T3Y) has a distinctive decapeptide core binding motif that exhibited variations between the P4-P6 residues that deviated from the classical conformation^39^. Although both AF2M-pred and AF3-pred accurately captured the variation at the P4 residue, they both encountered inaccuracy in predicting the subsequent binding conformations. These results suggested that AF-pred might be too loyal to canonical binding modes and not capable of predicting non-canonical binding modes.

**Figure 6.**
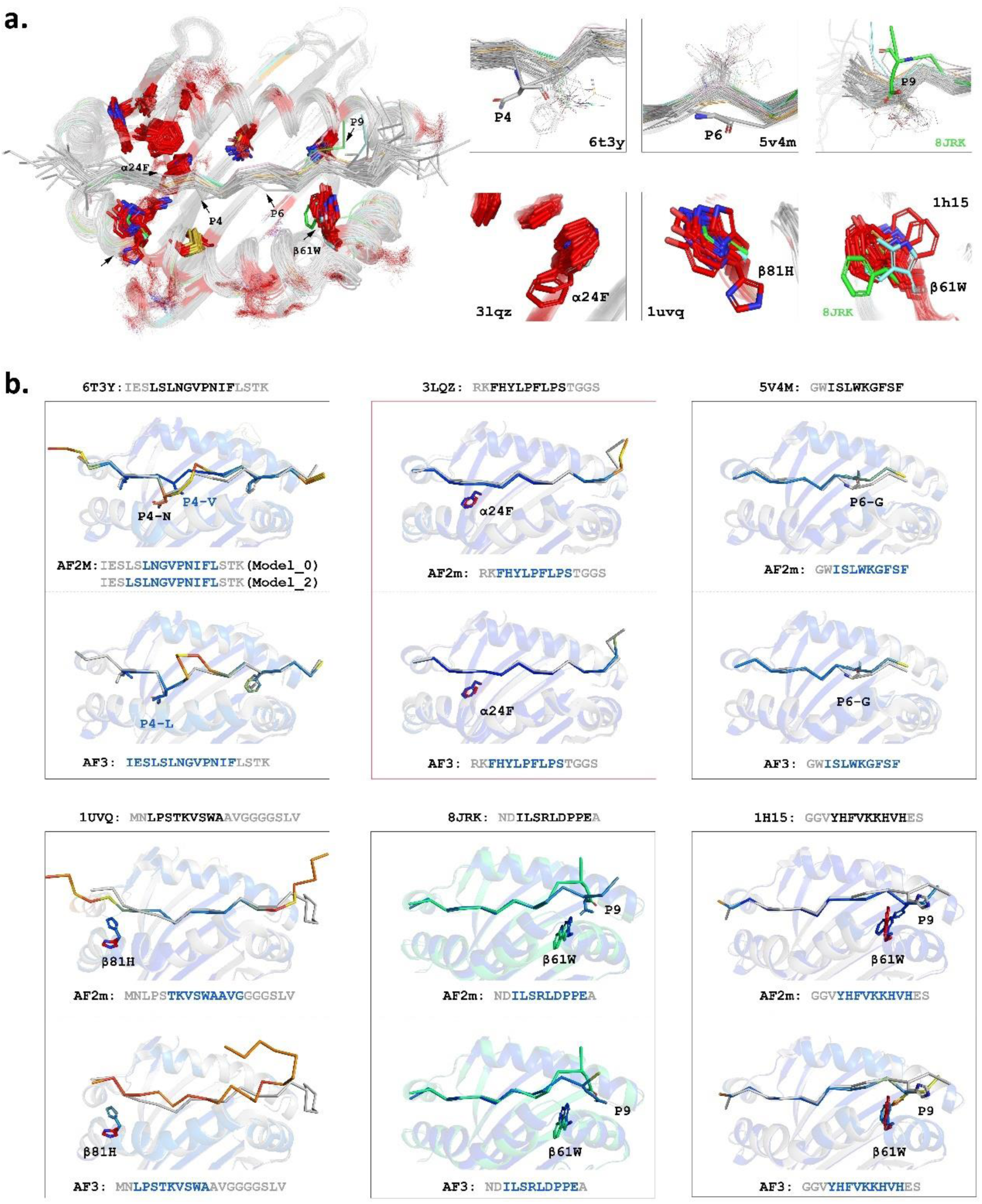
Analysis of AF’s inability to accurately predict non-canonical binding of peptides to MHC-II molecules. **(a).** Identification of structures with notable differences in peptide binding and MHC-II conserved residue conformations across resolved MHC-II complexes. **(b).** The AF predicted structures were not consistent with the crystal structures except the PDB entry 3LQZ.

## Discussion

Predicting MHC-II restricted epitopes is a challenging but very useful direction of applied research. Many tools for this application have been developed in the past decade, and they can be generally categorized into sequence-based and structure-based tools^12,13,23–25^. Although sequence-based tools show good performance in certain scenarios, they all require massive ligandome data for model training, making them not feasible for animal MHC-II with very limited ligandome data^27^. Structure-based prediction tools do not rely on massive ligandome data, but have strong generalization capabilities. So they have natural advantages in universal prediction capabilities for MHC-II restricted epitopes across alleles and species. Additionally, the conserved conformation of the 9aa-binding core region of MHC-II peptide ligands provides a clear standard for evaluating and interpreting the results of structure-based prediction results. AF has unprecedentedly improved the accuracy of protein structure prediction, providing the foundation for developing our new structure-based tool. In this study, we developed the AF-pred tool for predicting MHC-II restricted epitopes based on AF2M and AF3, and validated its prediction capability across species.

In the test with the human HLA-II peptidomic dataset, the best available sequence-based prediction tools NetMHCIIpan and MixMHC2pred showed higher accuracy than AF-pred. This result is not surprising: with heavy training with massive HLA-II ligandome data, NetMHCIIpan and MixMHC2pred are actually prediction tools specialized for human MHC-II. In the contrast, AF2M-pred and AF3-pred are generalized protein structure prediction tools designed for diverse application. Since optimizing the peptide binding characteristics of MHC-II or introducing peptide binding data for training and fine-tuning can significantly improve the performance of AF-pred^33–35^, we can certainly anticipate that iterative development of AF-pred finally lead to higher accuracy comparable with the best available sequence-based prediction tools.

However, for the prediction of MHC-II restricted epitopes across animal species, AF-pred demonstrated superiority over MixMHC2pred, which also claims to have cross-species prediction capability^31^. MixMHC2pred’s prediction capability comes from its training data, which includes peptide binding data from multiple species such as human, mouse, cow, and chicken MHC-II. Processing the correlation between the amino acid composition of MHC-II pockets and the corresponding anchor residues in the peptide ligands through specific neural network modules is the foundation of MixMHC2pred’s model. This enables the prediction of peptide binding characteristics of different MHC-II alleles based on pocket composition similarity. MixMHC2pred still requires massive non-human MHC-II ligandome data to train a workable model. Considering the differences in MHC-II sequences across species and the vast number of highly polymorphic alleles, accumulating massive ligandome data to complete the deep-learning model training is unrealistic in the near future.

In our study, AF-pred has demonstrated the capability of predicting MHC-II restricted epitopes without relying on massive ligandome data. This feature derived from its fundamental capability of protein structure prediction. This is an important inherent advantage for developing structure-based tools for predicting MHC-II restricted epitopes compatible for various animal species. In the meanwhile, the conserved peptide ligand binding mode to the MHC-II provide convenience for modeling and structure prediction.

Sequence-based tools have difficulties in predicting the affinity between binding peptide and MHC-I^35,40^. Structure-based tools can provide interaction force analysis between binding peptides and MHC-II, which can fundamentally solve this problem. Compared with the resolved crystal structures, both AF2M-pred and AF3-pred can accurately capture the conserved binding mode between MHC-II and peptide main chains and predict the subtle interactions between anchor residues and pockets, indicating their capability of predicting diverse MHC-II restricted epitopes. The thermal stabilities of the pBat_EF complexes formed by peptide PGDSIIRSMPEQTSEK and its derivative peptides are in high accordance with their predicted interaction forces (Fig. 5a). This indicated that AF-pred is capable of ranking the affinities of binding peptides by explaining their structural basis. However, there are still certain differences between the AF-pred output and the crystal structures, reflecting some limitations of the current tool.

First, AF-pred cannot accurately predict non-canonical peptide-MHC-II binding modes. Although the peptide binding mode of MHC-II is highly conserved, there are still certain exceptions that are categorized as non-canonical binding modes. The resolved pBat_EF-NDIL structure indicates that even the same peptide can bind MHC-II in different ways. However, so far there are not many crystal structure data representing non-canonical binding modes, and some newly resolved structures have not been included in the training dataset for AF-pred. This probably explains why AF-pred cannot make effective predictions for epitopes with non-canonical binding modes. But this also suggest that, involving more structures representing non-canoncical binding modes into the training of AF-pred may make up for the shortcoming in the future--we will certainly try this approach.

Second, AF-pred has preference on the most stable binding modes, so it may predict more molecular interactions than what really exist. Such preference could also be described as engaging as many as possible binding factors molecular interactions, but it may lead to false positive predictions. Motmaen, A. et al.^35^ found that AF2 tends to introduce non-binding peptides into the peptide-binding groove; but this problem can be effectively solved by adding a logistic regression layer at the top of the AF2 network to judge whether the peptide binds, as well as fine-tuning AF2 with binding and non-binding peptide training dataset. The determing factors of peptide-MHC-II binding include both favorable factors and restrictive factors. The above findings implied that AF-pred may identify redundant favorable factors and insufficient restrictive factors, leading to over-optimistic prediction results.

AF3-pred seem to idenfity restrictive factors in the structure better than AF2M-pred. According to the test results of a large number of HLA-II peptide ligands, AF3-pred demonstrated significantly higher accuracy than AF2M-pred. However, AF3-pred does not provide so much structural diversity as AF2M-pred in its output, probably because focusing on restrictive factors makes the prediction results more convergent. These differences may originate the replacement of the Pairformer module in AF3 compared to AF2M^38^, but our hypothesis needs further verification after AF3 is open-sourced. It is noteworthy that, AF-pred would be a very powerful tool for designing veterinary vaccines. SCMP scores for MHC-II binding and non-binding peptides, as well as the comparison of P/N values, have proven the rationality of the scoring logic and also provided a reference for setting confidence intervals for prediction results. SCMP>810 can be used as the criterion for high confidence prediction results. Despite some false-negative results, this criterion can maximize the positive prediction value (PPV) of epitope prediction, facilitating subsequent epitope immunogenicity verification work.

Compared with AF2M-pred, AF3-pred has advantages in prediction result rationality and computational resource consumption, making it more suitable for large-scale epitope screening. From the perspective of prediction accuracy, directly analyzing 9aa-peptide is better than the long peptide to determine the 9aa-binding core region. Therefore, the stepwise method can be used to screen all possible 9aa-binding core regions in a certain antigen as sources of MHC-II epitopes. The hotspot regions with intensive 9aa-binding core regions could be used for vaccine design. This method can avoid flanking regions’ interference on AF3-pred and provide fast prediction.

In summary, our research proves the feasibility of structure-based tools for predicting MHC-II restricted epitopes, as well as their advantage in generalization and interpretability across species. With tthe iterative development of AF-pred, the prediction accuracy can be further improved, greatly promoting the design and development of veterinary vaccines.

## Methods

### pMHC-II Dataset

The structures of the pMHC complex were downloaded from the RCSB Protein Data Bank (PDB)^41^, with an inclusion cutoff date set at 2024-04-05. The HLA-II binding data were extracted from the training sets generously provided by the NetMHCpan^40^ developers on their website (https://services.healthtech.dtu.dk/services/NetMHCIIpan-3.2/). The HLA-II binding dataset covers 10 pairs of HLA-II alleles, with 300 peptide ligands for each pair. Porcine and bovine MHC-II sequence data was procured from the IPD-MHC^42^ database (https://www.ebi.ac.uk/ipd/mhc/group), and their corresponding epitopes were derived from the P72 protein of African swine fever virus (ASFV). and the polypeptide of foot-and-mouth disease virus (FMDV). The bat MHC-II sequence data were obtained from the NCBI database (https://www.ncbi.nlm.nih.gov/), and peptide data were sourced from the S protein of COVID-19. The relevant MHC-II sequence and peptide sequence information mentioned above can be found in Supplementary Table 1.

### MHC-II restricted epitope Prediction Pipeline Based on AF

AF-pred is developed for the cross-species prediction of MHC-II restricted epitopes. Firstly, based on the input of paired MHC-II α/β chain sequences and ligand peptides, the structure of the pMHC-II complex was predicted using AF2M^37^ or AF3^38^. Consequently, by comparing with the resolved pMHC-II structure, the residues corresponding to the canonical 9aa-binding core region in MHC-II peptide ligands were automatically identified. The conformation of the 9aa-binding core region is highly conserved, and the perturbation between different pMHC-II structures does not exceed 2.5 Å. Hence, a pMHC-II structure (PDB ID: 1A6A) was selected, and the C-α atoms of its 9aa-binding core region were utilized as reference positions. After structural superposition, the correspondence between the C-α atoms in the predicted peptide and the reference C-α atoms is judged by a matrix algorithm based on the distances between the C-α atoms, and 2.5 Å is the acceptable threshold of correspondence. The number of corresponding C-α atoms reflects the probability of the predicted peptide to conform with the classical peptide binding conformation of MHC-II. Since peptide ligands may have two possible binding orientations in MHC-II peptide-binding groove: forward and reverse^22,31^, the distance matrix can determine the binding orientation of the predicted peptide by comparing the correspondence of C-α atoms in both forward and reverse directions. After determining the 9aa-binding core region and binding orientation, the flanking region (FR) of the predicted peptide will be subsequently determined.

The sum of the predicted local distance difference test (pLDDT) values of the corresponding C-α atoms in the predicted peptide is named as Sum_Core_Match_Result (SCMP). SCMP comprehensively reflects the prediction credibility, i.e. how the predicted peptide matches the conserved binding conformation in pMHC-II. Therefore, SCMP is used for ranking prediction results. To validate the prediction capability of AF-pred, we performed head-to-head comparison with two currently most successful sequence-based tools: netMHCIIpan-4.14 (https://services.healthtech.dtu.dk/services/NetMHCIIpan-4.1/) and mixMHC2pred-2.05 (http://mixmhc2pred.gfellerlab.org/).

### Purification of pMHC-II complex proteins

The extracellular regions of the MHC-II α-chain and β-chain were constructed into the pET-21a(+) vector (GenSmart™ 1.0 Gene Synthesis, GenScript Biotech Corporation). The ligand peptide was attached to the 5’ end of the β-chain via a Linker (SGGGGSLVPRGSGGGGS) using the MutExpress II Fast Mutagenesis Kit V2 (Vazyme, China).

The constructed plasmids were transformed into *E. coli* strain BL21(DE3) (Cat No.ZC206, ZOMANBIO, Beijing, China). After prokaryotic expression, the harvested inclusion bodies were dissolved in 6 M guanidine hydrochloride (JS0276, HongKong JiSiEnBei International Trade Co., Limited) at a final concentration of 30 mg/ml and stored at −20°C as described previously^43,44^. The solubilized inculsion bodies of MHC-II_α and peptide-MHC_β were slowly and dropwise added to the refolding buffer consisting of 100 mM Tris-HCl (pH 8.0), 2 mM EDTA, 30% W/V Glycerol, 0.9 mM oxidized glutathione and 3 mM reduced glutathione at a molar ratio of 1:2. After magnetically stirring process for 24 h at 4°C, the complexes were concentrated and switched into a buffer of 20 mM Tris-HCl (pH 8.0) and 50 mM NaCl. The complexes were further purified by gel filtration chromatography and anion-exchange chromatography (Superdex 200 16/60 column and Resource Q anion exchange column, GE Healthcare).

### Crystallization, data collection, and processing

3 peptides (PGDSFIIRSMPEQTS, GASFIIRSMPQET, NDILSRLDPPEA) were utilized to form crystals with bat-MHC-II_EF, respectively. The crystals were initially screened using the sitting-drop method with the Crystallization Screening Kit (Hampton Research). Purified pMHC-II complexes (referred as pBat_EF-PGDS, pBat_EF-GASF, and pBat_EF-NDIL, respectively) were diluted to the concentration of 2 or 4 mg/ml with molecular sieve buffer and then mixed with the crystallization screening kit buffer in a 1:1 volume ratio. pBat_EF-PGDS and pBat_EF-GASF crystals were grown as relatively thin slices in 0.2 M Calcium chloride dihydrate, 20% w/v Polyethylene glycol 3,350, pH 5.1. pBat_EF-NDIL crystal was grown as relatively thick slices in 0.2 M Ammonium formate, 20% w/v Polyethylene glycol 3,350, pH 6.6. Diffraction data were collected at Beamline BL18U1/BL19U of the Shanghai Synchrotron Radiation Facility using an R-AXIS IV++ imaging plate detector. The collected diffraction data were indexed and integrated using iMosflm^45^, and scaling and merging were performed using the CCP4i suite^46^.

The collected data was processed using CCP4i to determine the structure and calculate the phase via MOLREP^47^ and phasers^48^ (employing the molecular substitution method^49^ and using 1DLH as a structural template) to obtain the model and coordinates. After model building in COOT^50^ and several rounds of refinement in Refmac5^51^, further refinement was conducted using phenix refine^52^. Subsequently, the MolProbity^53^ tools were utilized in Phenix for assessing model quality^54^ (refer to Structure data table). Structure-correlation diagram analysis and plotting were performed using PyMOL (https://pymol.org/, Schrödinger, LLC).

## Supporting information

8JRJ

8JRK

## Reporting summary

Further information on research design is available in the Nature Portfolio Reporting Summary linked to this article.

## Data availability

The coordinate and structure factors presented in this article have been submitted to the Protein Data Bank (https://www.rcsb.org/) under accession numbers 8JRK, 8JRJ, 8JR4.

## Code availability

The code for the analysis of binding capacity between peptides and MHC-II has been uploaded to the Github website (https://github.com/limkong1992/pMHC2_CA_subpocket_count/blob/main/pMHC2_CA_subpocket_count.py).

Source code for the AF model, trained weights and an inference script are available under an open-source license at https://github.com/deepmind/alphafold.

The AF3 predictions were performed on the online AF3 server (https://golgi.sandbox.google.com/).

## Acknowledgments

This study was supported by the National Key Research and Development Program of China (2021YFD1800100), the National Natural Science foundation of China (32172871), The 2115 Talent Development Program of China Agricultural University.

## Contributions

S.W. carried out biological experiments and data analyses and picture mapping. L.K. and C.F. developed software and data analysis tools. D.H.,Z.T. and L.D. collated the experimental data. Sheng.W. and L.Z. conducted the data analysis. S.X. and H.Y. proofread the article. N.Z. provided the idea for the article and wrote the manuscript.

## Competing interests

The authors declare no competing interests.

## Supplementary Information

**Supplementary Table 1.MHC-II sequence and peptide sequence information**

**Supplementary Table 2. Predictive scores for 2000 peptides from the HLA-II datasets**

## Notes

https://services.healthtech.dtu.dk/services/NetMHCIIpan-3.2/

https://github.com/limkong1992/pMHC2_CA_subpocket_count/blob/main/pMHC2_CA_subpocket_count.py

### Competing Interest Statement

The authors have declared no competing interest.

